# Bacteria regulate choanoflagellate development with lipid activators, inhibitors, and synergists

**DOI:** 10.1101/052399

**Authors:** Arielle Woznica, Alexandra M. Cantley, Christine Beemelmanns, Elizaveta Freinkman, Jon Clardy, Nicole King

**Affiliations:** Howard Hughes Medical Institute and Department of Molecular and Cell Biology, University of California, Berkeley, California 94720, United States; Department of Biological Chemistry and Molecular Pharmacology, Harvard Medical School, 240 Longwood Avenue, Boston MA 02115, United States; Leibniz Institute for Natural Product Research and Infection Biology e.V., Hans-Knöll Institute (HKI), Beutenbergstrasse 11a, 07745 Jena, Germany.; Whitehead Institute for Biomedical Research, 9 Cambridge Center, Cambridge MA 02142, United States

## Abstract

In choanoflagellates, the closest living relatives of animals, multicellular “rosette” development is regulated by environmental bacteria. The simplicity of this evolutionarily-relevant interaction provides an opportunity to identify the molecules and regulatory logic underpinning bacterial regulation of development. We find that the rosette-inducing bacterium *Algoriphagus machipongonensis* produces three structurally divergent classes of bioactive lipids that, together, activate, enhance, and inhibit rosette development in the choanoflagellate *S. rosetta*. One class of molecules, the lysophosphatidylethanolamines (LPEs), elicits no response on its own, but synergizes with activating sulfonolipid rosette inducing factors (RIFs) to recapitulate the full bioactivity of live *Algoriphagus*. LPEs, while ubiquitous in bacteria and eukaryotes, have not previously been implicated in the regulation of a host-microbe interaction. This study reveals that multiple bacterially-produced lipids converge to activate, enhance, and inhibit multicellular development in a choanoflagellate.

**Significance Statement:** Bacterial symbionts profoundly influence the biology of their animal hosts, yet complex interactions between animals and their resident bacteria often make it challenging to characterize the molecules and mechanisms. Simple model systems can reveal fundamental processes underlying interactions between eukaryotes and their associated microbial communities, and provide insight into how bacteria regulate animal biology. In this study we isolate and characterize bacterial molecules that regulate multicellular development in the closest living relatives of animals, the choanoflagellate. We find that multiple bacterially-derived lipids converge to activate, enhance, and inhibit choanoflagellate multicellular development.

## Introduction

The foundational event in animal origins, the transition to multicellularity (1-3), occurred in oceans filled with diverse bacteria (4-7). There is a growing appreciation that specific bacteria direct diverse animal developmental processes, including light organ development in the Hawaiian bobtail squid and immune system development and maturation in organisms as diverse as cnidaria and mammals (8-20). However, the multicellularity of animals and the complex communities of bacteria with which they often interact hinder the complete characterization of many host-microbe dialogues.

Choanoflagellates, a group of microbial eukaryotes that are the closest living relatives of animals (21-24), promise to help illuminate the mechanisms by which bacteria influence animal development. As did cells in the first animals, choanoflagellates use a distinctive collar of actin-filled microvilli surrounding a flow-generating apical flagellum to capture bacteria as prey (25-27). Indeed, choanoflagellate-like cells likely formed the basis for the evolution of animal epithelial cells that today provide a selective barrier for mediating interactions with bacteria (27-29).

In many choanoflagellates, including *Salpingoeca rosetta*, a developmental program can be initiated such that single cells develop into multicellular “rosettes”. Importantly, rosette development does not occur through cell aggregation. Instead, as in the development of an animal from a zygote, rosettes develop from a single founding cell that undergoes serial rounds of oriented cell division, with the sister cells remaining stably adherent (Fig. 1). The orientation of the nascently divided cells around a central focus, the production of extracellular matrix, and the activity of a C-type lectin called Rosetteless, ultimately result in the formation of spherical, multicellular rosettes (30-32). Rosettes resemble morula stage embryos and the transition to multicellularity in *S. rosetta* evokes ancestral events that spawned the first animals (26, 27, 33).

**Fig. 1.**
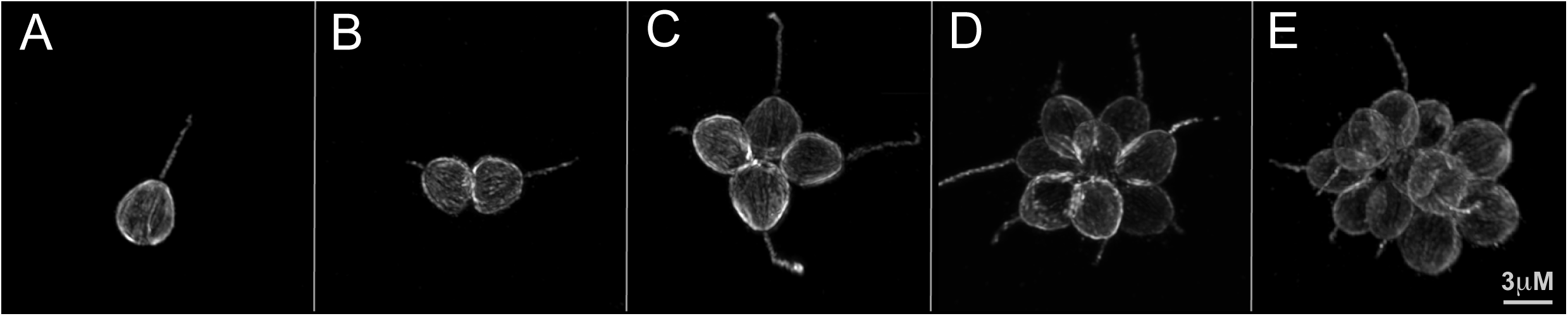
Stages of rosette development in *S. rosetta*. During rosette development, a single founding cell undergoes serial rounds of cell division, resulting in a structurally integrated rosette. Importantly, rosette development does not involve cell aggregation. Shown are a single cell (A), a pair of cells (B), a four-cell rosette (C), an eight-cell rosette (D) and a 16-cell rosette (E).

The initiation of rosette development was recently found to be induced by a co-isolated environmental bacterium, *Algoriphagus machipongonensis* (phylum Bacteroidetes; 34, 35). The ecological relevance of the *Algoriphagus* - *S. rosetta* interaction is evidenced by the co-existence of these organisms in nature (35), and the predator-prey relationship between choanoflagellates and bacteria (25, 36). Indeed, rosettes likely have a fitness advantage over single cells in some environments, as multicellular choanoflagellates are predicted to produce increased flux of water past each cell (37), and prey capture studies reveal that rosettes collect more bacterial prey/cell/unit time than do single cells (38). However, in other environments, rosette development would likely reduce fitness as rosettes have reduced motility relative to single cells. Therefore, we hypothesize that choanoflagellates utilize bacterially-produced molecules to identify environments in which rosette development might provide a fitness advantage.

The simplicity of the interaction between *S. rosetta* and *A. machipongonensis* (hereafter, *‘Algoriphagus’*), in which both members can be cultured together or independently, offers a biochemically tractable model for investigating the molecular bases of bacterial-eukaryotic interactions. Using rosette development as a bioassay, the first rosette-inducing molecule, <u>R</u>osette <u>I</u>nducing <u>F</u>actor-1 (RIF-1), was isolated from *Algoriphagus*. The observation that RIF-1 fails to fully recapitulate the bioactivity of the live bacterium (Figs. 2A and C), raises the possibility that additional molecules might be required (35). To gain a more complete understanding of the molecules and regulatory logic by which bacteria regulate rosette development, we set out to identify the minimal suite of *Algoriphagus* molecules that are necessary and sufficient to regulate *S. rosetta* rosette development.

**Fig. 2.**
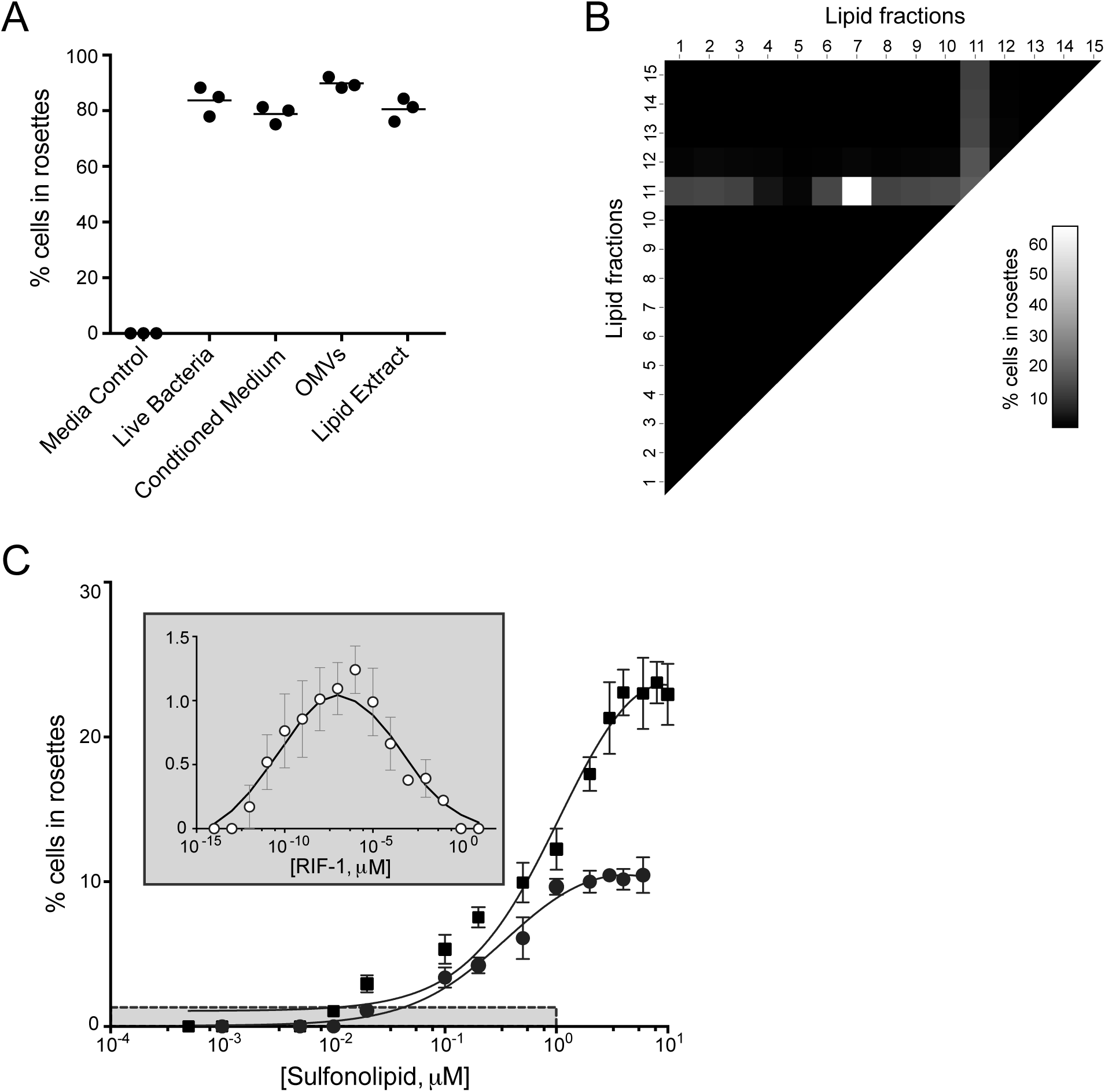
Maximal rosette development requires lipid co-factor interactions. (A) When treated with media that lacks necessary bacterial signals (Media Control), *S. rosetta* does not produce rosettes. In contrast, when treated with live *Algoriphagus, Algoriphagus* conditioned media, *Algoriphagus* OMVs, or *Algoriphagus* bulk lipid extract, rosettes develop at maximal (~90% cells in rosettes) or near-maximal levels. **(B)** A heat map depicts the rosette-inducing activity of *Algoriphagus* lipid fractions used to treat SrEpac, either in isolation or in combination, at a final lipid concentration of 2 μg/mL. Sulfonolipid-enriched fraction 11 was the only fraction sufficient to induce rosette development when tested alone (30% of cells in rosettes). Tests of each of the lipid fractions in combination revealed previously unidentified inhibitory and enhancing activity. Fractions 4 and 5 decreased rosette development (to 12% and 8%, respectively) in fraction 11-treated cells, whereas fraction 7 increased rosette development to 65%. **(C)**The RIF mix (solid square) and purified RIF-2 (solid circle) induced rosette development at μM concentrations. Inset: RIF-1 (open circle) is active at fM to nM concentrations, but induces 10-fold lower levels of rosette development than RIF-2. Grey box in main graph indicates the range of concentrations at which RIF-1 is active, and the range of its rosette-inducing activity. Rosette development was quantified 24 hours after induction. Minor ticks on X-axis are log spaced.

## Results

### A newly identified sulfonolipid activates the rosette development pathway

To identify the minimal set of *Algoriphagus* molecules required for full rosette induction, we used a bioassay based on a co-culture of *S. rosetta* with the non-rosette inducing prey bacterium *Echinocola pacifica* (see SI Materials and Methods). This culture, called ‘SrEpac’ (for *S. rosetta* + *E. pacifica*; 39), reproducibly yields high percentages of cells in rosettes (>80%) in response to live *Algoriphagus*, *Algoriphagus* outer membrane vesicles (OMVs; see SI Text) isolated from conditioned medium, and *Algoriphagus* bulk lipid extracts (Fig. 2A; SI Appendix, Fig. S1). In addition, incubation of SrEpac with the only previously known <u>R</u>osette <u>I</u>nducing <u>F</u>actor, the sulfonolipid RIF-1, results in low but reproducible levels of rosette development (~1.5% of cells in rosettes; Fig. 2C), consistent with previous results using a different *S. rosetta* culture (35).

Because *Algoriphagus* bulk lipid extracts elicit the same rosette development response as live bacteria (Fig. 2A), we began by fractionating a bulk extraction of *Algoriphagus* lipids by reversed-phase high performance liquid chromatography (HPLC) and testing the resulting 15 lipid fractions in SrEpac (see SI Materials and Methods). Only fraction 11 was sufficient to induce rosette development, whereas all other lipid fractions lacked rosette-inducing activity at all concentrations tested (Fig. 2B). To further separate and isolate the active molecules in fraction 11, we performed a subsequent round of reversed-phase HPLC and tested the resulting sub-fractions for activity in SrEpac. The rosette-inducing activity tracked with one sub-fraction (hereafter, “RIF mix”) that induced rosette development in 23.5% of cells (Fig. 2C; SI Appendix, Fig. S2). Structural analysis by NMR, high resolution mass spectrometry (HRMS), and tandem mass spectrometry (MSMS) revealed that the RIF mix contained RIF-1 and two structurally related but previously uncharacterized sulfonolipids with approximate molecular weights of 605 Da and 593 Da (SI Appendix, Figs. S2 −17). Sulfonolipids are a largely uncharacterized class of molecules that are structurally similar to sphingolipids, a diverse group of molecules based on sphingoid bases that play structural roles in cell membranes and important non-structural roles in signal transduction (40). Although sulfonolipids have been reported to contribute to the gliding motility of Bacteroidetes bacteria (41, 42), almost nothing is known regarding their potential roles as signaling molecules.

Additional activity-guided fractionation by HPLC allowed us to isolate pure samples of RIF-1 (35, 43) and of the 605 Da sulfonolipid. Purified RIF-1 induced maximal (~1.5%) rosette development at femtomolar to nanomolar concentrations (Fig. 2C, inset). In contrast, the purified 605 Da sulfonolipid (hereafter “RIF-2”) elicited 7-fold higher levels of rosette development (10.5% of cells in rosettes; Fig. 2C) than RIF-1, although at micromolar concentrations. The planar structure of RIF-2 (Fig. 3A) was determined by one and two-dimensional NMR (see SI Appendix Table S1, Figs. S3-17), and was found to closely resemble RIF-1, with the exception of slight structural variations of the capnoid base, which contains a double bond at C-4 and a hydroxyl group at C-6.

**Fig. 3.**
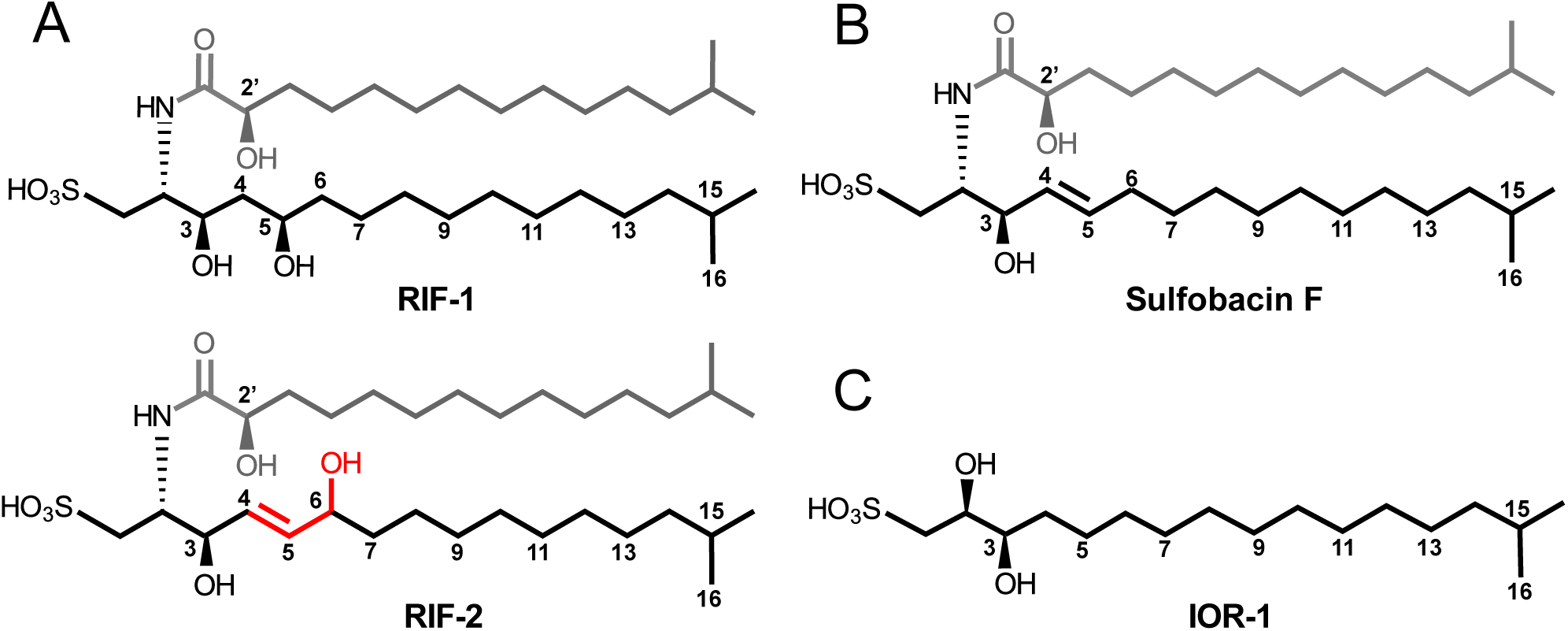
Structural similarities and differences among RIFs, an inactive sulfonolipid, and the inhibitory capnine IOR-1. **(A)** The 3D structure of RIF-1 (43), compared to the proposed molecular structure of RIF-2, and **(B)** the structure of an inactive *Algoriphagus* sulfonolipid, Sulfobacin F (43). Shared features of *Algoriphagus* sulfonolipids include a fatty acid chain (shown in grey), and a capnoid base (shown in black). Distinguishing features between RIF-1 and RIF-2 (highlighted in red) include a double bond at position 4, and a hydroxyl group at position 6. The tight structure-activity relationships of RIF-1 and RIF-2 suggest a restricted set of interactions between these molecules and a binding target. No features are shared by RIF-1 and RIF-2 to the exclusion of Sulfobacin F. **(C)** The IOR-1 capnine antagonizes the rosette inducing activity of RIFs. Like the capnoid base of RIF sulfonolipids, IOR-1 is composed of a sulfonic acid head group and a branched chain containing two-OH groups. Furthermore, the carbon chain length and branching pattern of IOR-1 matches the capnoid base in RIF-1 and −2. The similarities between IOR-1 and the RIFs raise the possibility that IOR-1 competitively inhibits RIF binding to a target receptor.

The remaining 593 Da sulfonolipid in the RIF mix is produced by *Algoriphagus* at low levels (approximately 1/5^th^ the amount of RIF-2) and elutes closely to RIF-2 during fractionation. Although HRMS and HRMSMS data suggest that this molecule is a sulfonolipid similar to RIF-1, low levels of production and co-elution with RIF-2 have prevented us from fully isolating and characterizing the activity of the 593 Da sulfonolipid (SI Appendix, Figs. S2, S6). However, because the combination of RIF-2 and the 593 Da sulfonolipid induced rosettes at levels indistinguishable from those of RIF-2 alone (SI Appendix, Fig. S18), we infer that the rosette-inducing activity of the RIF mix is largely the product of RIF-2. Nonetheless, we note that the maximal level of rosette development induced by the RIF mix (Fig. 2C) is greater than the sum of purified RIF-1 + RIF-2, for reasons that we do not yet understand.

The discovery of RIF-2 revealed that RIF-1 is not the sole *Algoriphagus* determinant of *S. rosetta* rosette development. However, even the RIF mix, which contains both RIF-1 and RIF-2, failed to recapitulate the full level of rosette induction elicited by either intact *Algoriphagus* or *Algoriphagus* bulk lipid extract. Therefore, we hypothesized that additional molecular cues are required to fully potentiate the rosette-inducing activities of RIF-1 and −2.

### Lipid cofactors inhibit and enhance RIF activity

To identify potential cofactors of the RIFs, we mixed each of the 15 *Algoriphagus* lipid fractions in pairwise combinations and tested the mixtures at several concentrations in SrEpac (Fig. 2B; SI Appendix, Materials and Methods). We observed two types of cofactor activity: enhancing activity in fraction 7 and, unexpectedly, inhibitory activity in fractions 4 and 5. Importantly, the activities of these cofactor-containing fractions were only evident when tested in combination with fraction 11, which contained both RIF-1 and RIF-2.

The inhibitory activity observed in fractions 4 and 5 is the first example of a compound(s) – either isolated from *Algoriphagus* or commercially available – that specifically reduces levels of rosette development at concentrations that do not otherwise inhibit growth (SI Appendix, Table S2). Therefore, we used bioactivity-guided fractionation in the presence of RIF-2 to determine the molecular basis for inhibition. HRMS and NMR experiments, together with total synthesis (44), allowed us to propose the absolute structure for the 351 Da molecule (hereafter referred to as Inhibitor of Rosettes-1, “IOR-1” Fig. 3C). Comprehensive methods detailing IOR-1 isolation and structure determination, along with dose-response curves of IOR-1 in the presence of the RIF mix and RIF-2, are described in (44).

Nanomolar concentrations of IOR-1 completely inhibits the ability of RIF-2 to induce rosette development, and reduces rosette development in the presence of the RIF mix (SI Appendix, Fig. S19). IOR-1 is a capnine lipid that resembles the capnoid backbone of *Algoriphagus* RIFs (Figs. 3A and C). Thus, we hypothesize that IOR-1 antagonizes rosette development by competitively binding a RIF-2 target receptor. Because the RIF-mix induces low levels of rosette formation in the presence of IOR-1, we infer that the combined effects of the RIFs are sufficient to partially overcome the presence IOR-1.

In contrast to the inhibitory activity associated with IOR-1, the *Algoriphagus* lipid fraction 7 greatly enhanced rosette development when used in combination with the RIF-containing fraction 11 (Fig. 2B). Notably, fraction 7 did not contain any sulfonolipids, the only class of molecules previously known to regulate rosette development. After separating the components of fraction 7 by HPLC, we treated SrEpac with each subfraction in combination with the RIF mix and quantified the level of rosette development. The subfractions that enhanced rosette development in the presence of the RIF mix contained one or both of two lysophosphatidylethanolamines (LPEs) with molecular weights of 451 Da and 465 Da (hereafter referred to as LPE 451 and LPE 465, respectively; Fig. 4A).

**Fig. 4.**
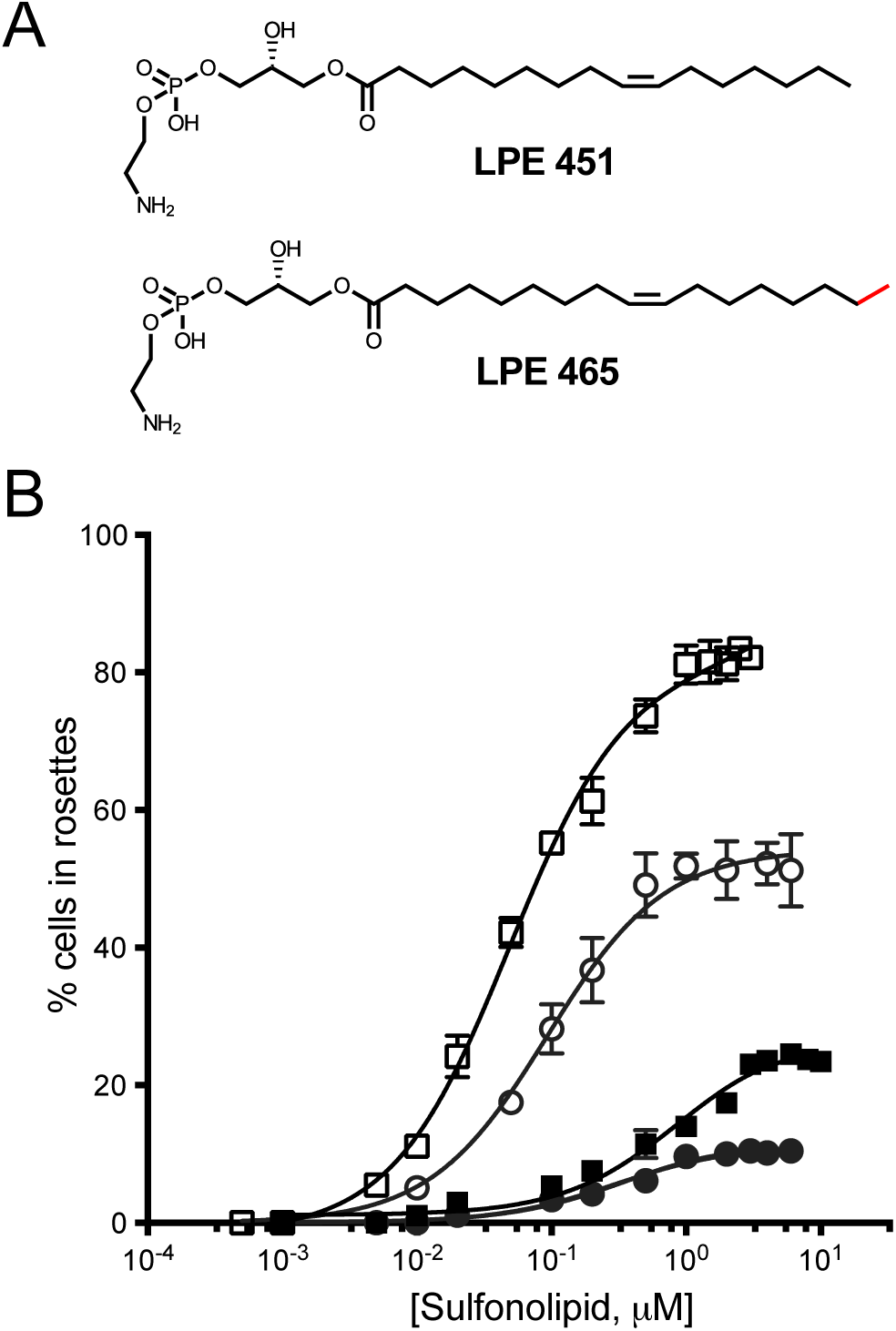
LPEs synergize with RIFs to enhance rosette development. **(A)** The structures of LPE 451 and LPE 465 as determined by NMR and tandem mass spectrometry. LPE 451 and LPE 465 differ by only one methyl group (highlighted in red). (**B**) The addition of 2 μM LPE mix increases the maximal percentage of cells in rosettes in RIF-2-treated SrEpac from 10.5% (solid circle) to 52% (open circle) and the maximal inducing activity of the RIF mix from 23.5% (solid square) to 82% (open square) of cells in rosettes. Minor ticks on X-axis are log spaced.

As this class of molecules is well known, literature precedence allowed us to confirm the core LPE structure by NMR and tandem mass spectrometry (Fig. 4A; SI Appendix, Figs. S20-27). We performed an olefin metathesis on the most active LPE fractions (45) to determine that the major species present (in both LPE 451 and LPE 465) contains a double bond between carbons 9 and 10, which is common for fatty acid chains of this length (SI Appendix, Fig. S28). Due to the difficulties associated with purifying these types of molecules, we were unable to completely exclude other LPE isoforms (which can differ in double bond location or position on the glycerol backbone); however, multiple iterations of bioassay-guided fractionation consistently yielded a fraction from the purification process (hereafter, the “LPE mix”) in which 98% of the fraction was composed of LPEs 465 and 451, with the remaining 2% of the sub-fraction containing trace amounts of other structurally related LPE analogs. Importantly, no commercially available LPEs tested in combination with the RIF mix either activated or enhanced rosette development (SI Appendix, Table S2). Therefore, we infer that LPE 451, LPE 465, or both, are responsible for the synergistic RIF-enhancing activity of the LPE mix. Furthermore, as with the RIFs (43) and IOR-1 (44), it appears that the enhancing activity of the LPEs results from a highly specific structure-activity relationship.

LPEs belong to a large and diverse class of deacylated phospholipids, called lysophospholipids, that include structural components of cellular membranes as well as biologically active lipid mediators (46, 47). While LPEs are found in most bacterial and eukaryotic cell membranes and present in somewhat elevated concentrations in many marine and estuarine bacteria (48), little is known about how and in what contexts LPEs might act as signaling molecules (47, 49).

To characterize how LPEs regulate rosette development, we started by investigating the concentrations at which the LPE mix displayed maximal enhancing activity. In contrast with the 10.5% of cells in rosettes induced by 2 μM RIF-2 alone, treatment of SrEpac with 2μM RIF-2 and micromolar concentrations of the LPE mix increased rosette development five-fold to 53% (Fig. 4B; SI Appendix, Fig. S19). Furthermore, maximal levels of rosette development elicited by the RIF mix + the LPE mix matched those induced by the *Algoriphagus* lipid extract (Fig. 2A; Fig. 4B).

Finally, we observed that LPEs also influence RIF potency. In bioassays in which the concentration of the LPE mix was held stable at 2 μM and the RIF mix or RIF-2 was titrated, the sensitivity of *S. rosetta* to the RIFs increased such that 25-fold less RIF mix and 3-fold less RIF-2 was required to achieve half-maximal induction (Fig. 4B). These results reveal that the rosette inducing activity of *Algoriphagus* can be largely recapitulated with specific representatives from just two different classes of lipids: sulfonolipids and LPEs.

### LPEs promote a previously unidentified maturation step in rosette development

Rosettes induced by live *Algoriphagus* bacteria or *Algoriphagus* OMVs, lipid-rich vesicles that fully recapitulate the inducing activity of live bacteria, are remarkably resistant to shear and can range in size from 4 cells, the minimum number of cells required to confirm the organized polarity of a rosette (see SI Materials and Methods), to as many as 50 cells. Because the rosette-inducing activity of OMVs is stable, highly reproducible, and equivalent to that of live bacteria, we used it as a positive control for the study of rosette cell number. Within just 22 hours after treatment, OMV*-*induced rosettes were resistant to shear introduced by pipetting, and the median cell number per rosette was 8 cells, although some grew to as large as 16 cells/rosette (Fig. 5A). In contrast, treatment with purified RIF-2 resulted in rosettes that were sensitive to mechanical disruption; after pipetting, the median cell number per rosette was significantly smaller (4 cells/rosette) than that induced by *Algoriphagus* OMVs (8 cells/rosette; Fig. 5A). Furthermore, the size frequency distribution for RIF-2-induced rosettes was restricted to small rosettes, ranging from the minimum size of 4 cells up to 8 cells, compared to *Algoriphagus*- and OMV-induced cultures in which larger rosettes of 10-16 cells were frequently observed. Because the combinatorial activity of RIF-2 + LPE mix resulted in elevated percentages of cells in rosettes, we hypothesized that LPEs might promote rosette stability and therefore protect larger rosettes when exposed to shear. Indeed, the median cell number (7 cells/rosette) and size frequency distribution of SrEpac induced by RIF-2 + LPE mix was statistically indistinguishable from OMV-induced cultures (Fig. 5A).

**Fig. 5.**
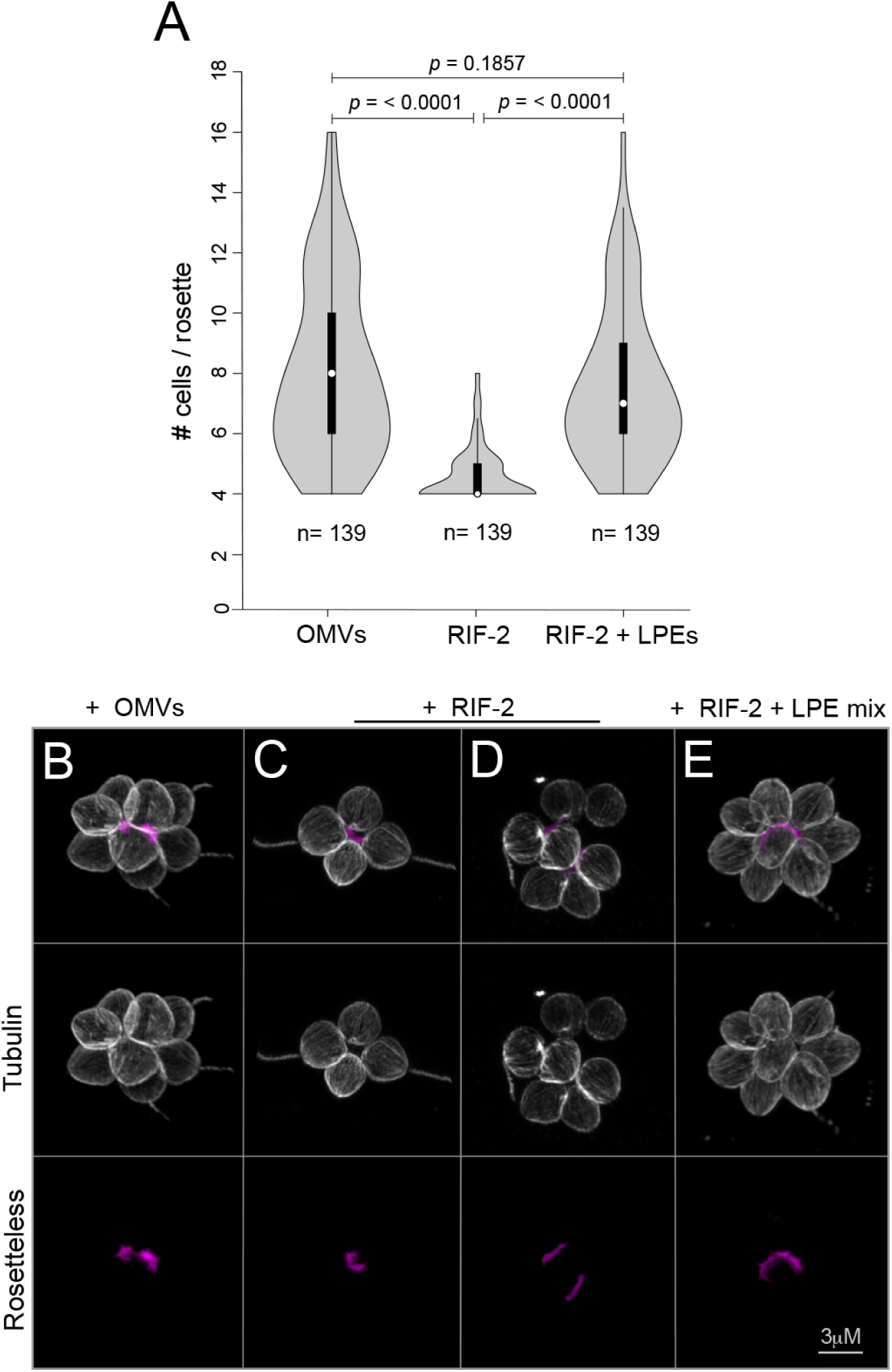
LPEs promote proper rosette development and maturation. (**A**) Frequency distribution of rosette size in SrEpac incubated with OMVs, RIF-2, and RIF-2 + LPE mix after exposure to shear stress by pipetting. Rosettes induced by RIF-2 alone contained fewer cells on average and reached a smaller maximal size than rosettes induced with *Algoriphagus* OMVs. The addition of the LPE mix to RIF-2 increased the median rosette size and frequency distribution to levels that recapitulated induction by OMVs. Rosette size was assessed 22 hours after induction (n=139 for each condition). Violin box plots show the median cell number (white circle), 75% quartile (thick line), and range excluding outliers (thin line). Surrounding the box plot is a kernel density trace, plotted symmetrically to show rosette size frequency distribution. P values (unpaired t-tests) were calculated using GraphPad Prism v6 for Mac, GraphPad Software, La Jolla, CA, USA. **(B-E)** Rosette morphology, cell packing, and localization of Rosetteless protein (a marker of rosette development) in rosettes induced by **(B)** OMVs, **(C and D)** RIF-2 alone, and **(E)** RIF-2 + LPEs. (**B**) Cells in OMV-induced rosettes express Rosetteless and are tightly packed. Anti-tubulin antibodies (white) highlight the cell body and anti-Rosetteless antibodies (magenta) stain Rosetteless protein in the center of rosettes (Levin et al., 2014). **(C)** 4-celled rosettes induced by RIF-2 are tightly packed, whereas larger rosettes induced by RIF-2 alone **(D)** appear disorganized, with cells spaced farther apart. **(E)** Rosettes induced by RIF-2 + LPE mix are large and closely packed, and phenocopy rosettes induced by OMVs. All rosettes were fixed 22 hours after treatment.

The hypothesis that RIF-2 induced rosettes exhibit less structural integrity than rosettes induced by either OMVs or RIF-2 + LPEs was supported by observations made using high-resolution microscopy (Figs. 5B-E). Cells in OMV-induced rosettes were tightly packed and properly localized a specific marker of rosette development, the C-type lectin protein Rosetteless (32), to the extracellular matrix-rich center of the rosette (Fig. 5B). While 4-celled rosettes induced by RIF-2 alone showed close cell packing, cells in all larger rosettes induced by RIF-2 (e.g. those with 5-7 cells/rosette) were spaced farther apart than those in OMV-induced rosettes of equivalent size (Fig. 5D). Despite a ‘loose’ morphology, RIF-2-induced rosettes secreted Rosetteless protein, demonstrating that they had properly initiated rosette development. Importantly, induction with RIF-2 + the LPE mix restored a robust rosette morphology, with the cells tightly packed together, phenocopying OMV-induced rosettes. Thus, although RIFs alone are sufficient to initiate rosette development, LPEs promote structural stability during rosette development, and thereby facilitate rosette maturation (Fig. 6).

**Fig. 6.**
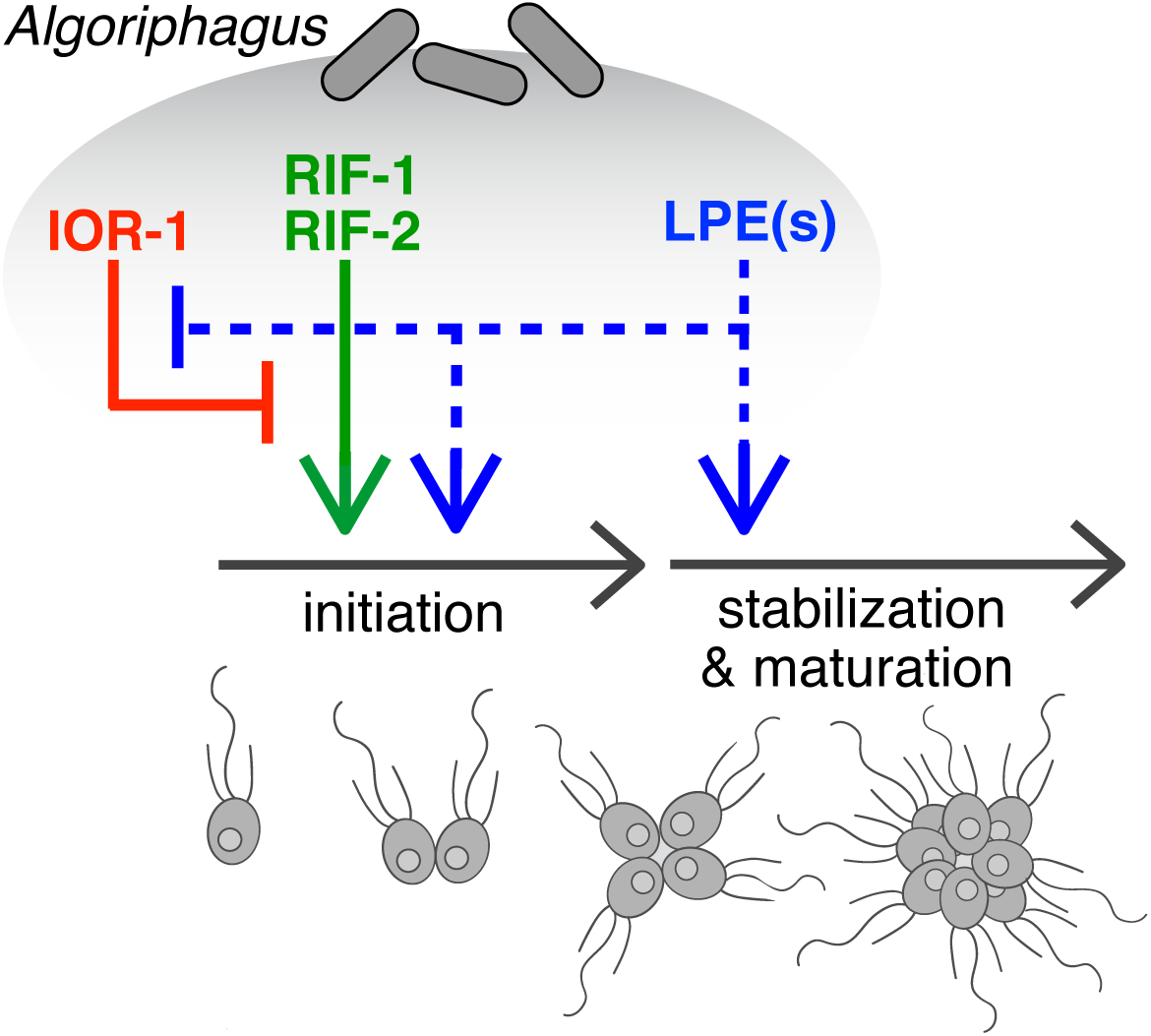
Multiple bacterial inputs regulate rosette development in *S. rosetta*. *Algoriphagus* produces three chemically distinct classes of lipids – sulfonolipids, LPEs, and a capnine – that interact to alternately induce, enhance, or inhibit rosette development in *S. rosetta*. The sulfonolipids RIF-1 and RIF-2 are sufficient to initiate rosette development in *S. rosetta*, although rosettes induced by RIFs alone are restricted in size, potentially because of their sensitivity to shear. Complete rosette maturation requires the synergistic activities of RIFs and LPEs. Although LPEs have no detectable activity on their own, they enhance RIF activity and facilitate the growth of larger and more structurally stable rosettes, perhaps by regulating downstream pathways important for rosette maturation. While the molecular mechanisms by which LPEs regulate rosette development are unknown (indicated by dotted lines), multiple lines of evidence (see main text) suggest that LPEs act both to promote the initiation of rosette development and, separately, to promote the subsequent maturation of rosettes. *Algoriphagus* also produces the inhibitory molecule IOR-1, which inhibits the rosette-inducing activity of RIFs (44). Importantly, when *S. rosetta* is exposed simultaneously to RIFs and the synergistic LPEs, mature rosettes develop even in the presence of IOR-1.

## Discussion

Animals rely on bacteria for everything from proper metabolism to the stimulation of immune system development to the regulation of gut morphogenesis (19, 50). Bacterial cues even direct major life history transitions in animals, with many marine invertebrates producing motile larvae that will not settle and undergo morphogenesis until they encounter the appropriate environmental bacteria (51). In one of the most dramatic examples of cross-talk between bacteria and an animal, *Vibrio fischeri* bacteria are recruited into crypts in the juvenile Hawaiian bobtail squid, where the bacteria then trigger post-embryonic morphogenesis of the “light organ” (20). The widespread phylogenetic distribution of bacterially-regulated developmental processes in animals suggests that such interactions may have been pivotal during the origin and early evolution of animals (19, 52).

As the number of animal developmental processes influenced by bacteria grows, detailed molecular characterization of the relevant bacterially-produced cues promises to reveal the regulatory logic underlying host-microbe interactions. Through the study of rosette development in a close relative of animals, *S. rosetta*, we have found three classes of structurally distinct lipids produced by *Algoriphagus* that are interpreted by *S. rosetta* as activators, synergistic enhancers, and inhibitors of development (Fig. 6). When tested alone, activating RIFs elicit relatively low levels of rosette development and the synergistic LPEs have no detectable activity. However, when used in combination, the activating RIFs + synergizing LPEs induce levels of rosette development in *S. rosetta* that recapitulate those induced by live *Algoriphagus* (Figs. 2 and 4). Moreover, while the *Algoriphagus* capnine IOR-1 is a potent antagonist of the RIFs (44), the synergistic activities of the RIFs and LPEs overcome the inhibitory activities of IOR-1, potentially explaining why endogenous IOR-1 does not prevent robust rosette induction.

We hypothesize that the reliance of *S. rosetta* on multiple inputs from *Algoriphagus* prevents the developmental switch to rosette development under suboptimal conditions. The commitment to rosette development requires a trade-off; rosette development is a lengthy process and while rosettes are potentially more efficient than single cells in the capture of planktonic bacteria, they are poor swimmers (53) and therefore likely to be less effective at dispersal and escape from certain predators (e.g. amoebae). Moreover, the aquatic world in which choanoflagellates live is patchy (54), with the diversity and density of bacteria dramatically varying between local microenvironments. In animals, the integration of multiple signals is fundamental to the robustness of many developmental decisions, including the establishment of the body axis during early embryogenesis (55-58), and the progressive specification of cell fates (59-61). Likewise, the multi-input regulatory module that controls *S. rosetta* development may act to ensure that rosette development is not initiated under the wrong environmental conditions or in response to the wrong bacterial cues.

The integration of multiple bacterial inputs is also essential for proper animal development in two well-studied host-microbe models. In the Hawaiian bobtail squid, two molecules (LPS and TCT) produced by *Vibrio fischeri* act synergistically to trigger light organ maturation (10), and in mice, several bacterial cell wall molecules (LPS, PGN, and polysaccharide A) together shape the development of the immune system of the gut (12, 13, 62). The finding that rosette development in *S. rosetta* requires the integration of a network of bacterial lipids extends this phenomenon to the closest living relatives of animals. Ultimately, as the molecular underpinnings of more host-microbe interactions are fully elucidated, the mechanisms by which bacteria influence their animal hosts may be found to be as intricate and complex as those regulating animal development, with microbial communities providing cocktails of activating, enhancing, and inhibitory cues.

## Materials and Methods

*Detailed methods are provided in **SI Appendix, Materials and Methods***.

**Choanoflagellate husbandry and rosette development bioassays**. SrEpac was propagated in 5% Sea Water Complete media. For all rosette development bioassays, cultures of SrEpac were diluted to a density of ~10^4^-10^5^ cells/mL prior to treatment with bacterial fractions and purified lipids.

**Quantifying rosette development**. To determine the percentage of cells in rosettes, the relative numbers of single cells and cells within rosettes were scored until 500 total cells had been counted per technical replicate. To quantify rosette size, the number of cells in each rosette was counted for each group of four or more cells with organized polarity relative to a central focus (with each cell oriented with the apical flagellum pointing outward) after exposure to shear. Rosette integrity was also characterized by immunofluorescence microscopy.

**Fractionation, isolation, and characterization of *Algoriphagus* lipids**. Ethyl acetate extracted *Algoriphagus* lipids were separated by reversed phase HPLC and tested for induction of rosette development in SrEpac. Subsequent rounds of activity guided fractionation led to preparations that were sufficiently pure for characterization by LC-MS/MS and NMR.

## Acknowledgements

We thank M. Abedin, D. Booth, B. Davidson, and J. Rawls for critical reading of the manuscript, R. Alegado for early discussions, L. Wetzel for help with ‘violin’ plots, and J. Wang at the Harvard Small Molecule Mass Spec Facility for help with high resolution mass spec. NK is a Senior Fellow in the Integrated Microbial Biodiversity Program of the Canadian Institute for Advanced Research.

